# Ligand-dependent G protein dynamics underlying opioid signaling efficacy

**DOI:** 10.1101/2025.06.30.662233

**Authors:** Jonathan Deutsch, Daniel Hilger, John Janetzko, Charles M. Schroeder, Steven Chu, Brian K. Kobilka, Rabindra V. Shivnaraine

## Abstract

Activation of heterotrimeric G proteins by G protein-coupled receptors (GPCRs) requires large-scale opening of the Gα α-helical domain (AHD) to expose the nucleotide-binding site and facilitate GDP–GTP exchange. While orthosteric ligands are known to modulate GPCR conformation and signaling efficacy, how these effects propagate to the G protein itself remains unclear. Using single-molecule fluorescence resonance energy transfer (smFRET) imaging, we monitored AHD motions in G_i_ proteins coupled to the μ-opioid receptor (μOR) across a spectrum of ligand- and nucleotide-bound states. We find that receptor ligands differentially modulate these dynamics from over 70 Å away, with higher-efficacy agonists more effectively promoting transitions to an open, low-nucleotide-affinity conformation. These data also capture transient μOR-G_i_ intermediates during nucleotide binding and suggest that μ-opioid ligand efficacy arises in part from allosteric control over G protein conformational equilibria that kinetically gate activation.

## Main

Full and partial agonists have distinct therapeutic applications for targeting opioid receptors, with full agonists used in pain management and partial agonists in treatment of opioid use disorder. μ-opioid receptors (μORs) are GPCRs that mediate analgesia and reward responses to common opioid drugs by catalyzing guanine nucleotide exchange in heterotrimeric G_i_ proteins, composed of Gα, Gβ, and Gγ subunits^1^. When activated by ligands, μOR and other class A GPCRs couple to inactive, GDP-bound G proteins at their intracellular surface (Fig. 1a). This interaction induces conformational changes in the G protein that promote GDP release and GTP binding, activating the G protein and triggering downstream signaling cascades which effectuate the physiological response^2^. Recent biophysical evidence indicates that ligands modulate GPCR signaling by stabilizing distinct receptor conformational states that vary in their G protein coupling efficiency^3–9^. However, the extent to which these ligand-dependent states propagate to the coupled G protein remains unclear, and if they do, whether these altered G protein conformational states also affect signaling outcomes in the cell.

**Fig 1.**
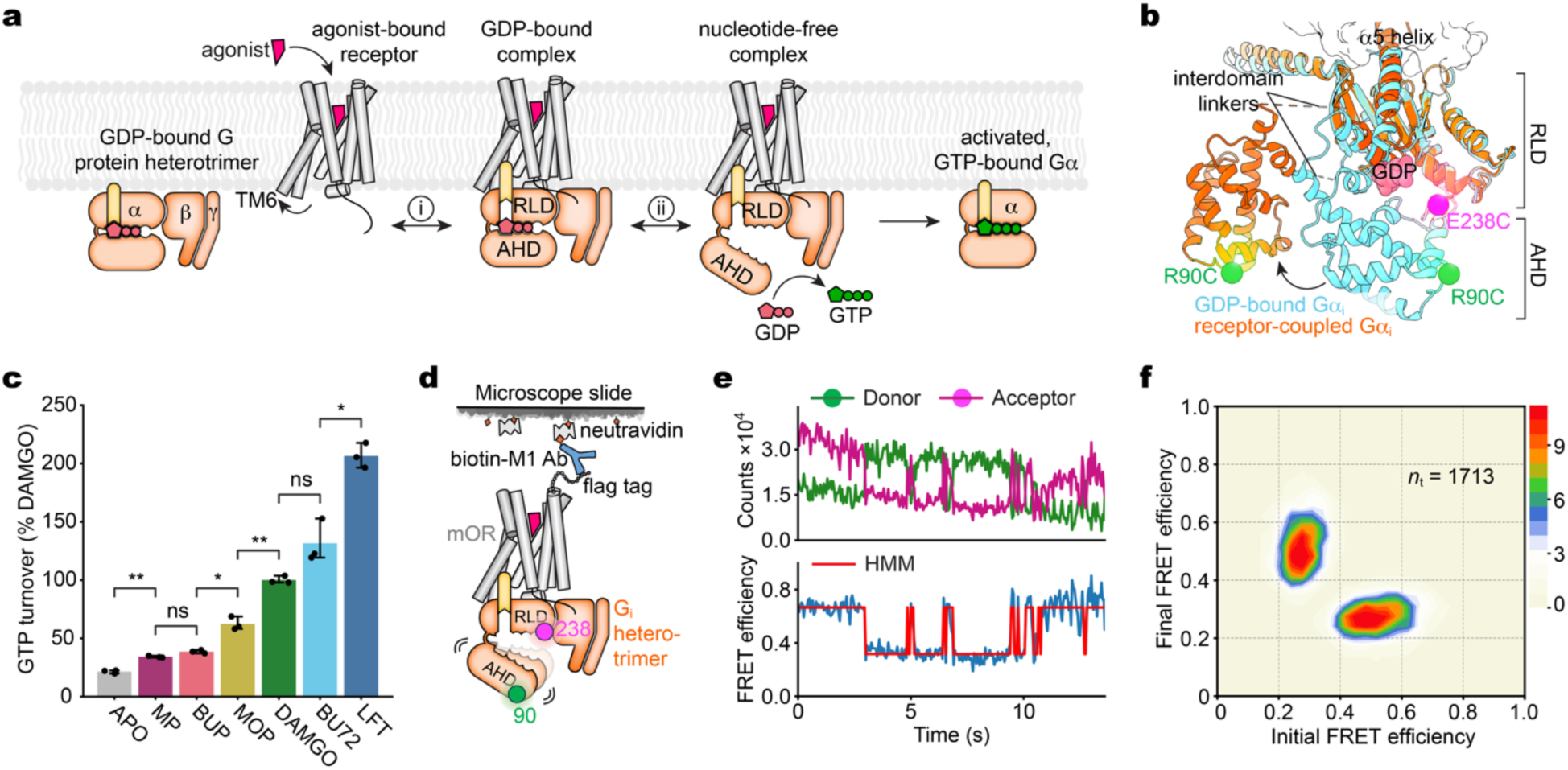
Intrinsic G protein conformational dynamics facilitate μOR-G_i_ activation. **a**, Agonist binding to a class A GPCR promotes G protein coupling and activation via GDP-GTP exchange. The activation rate can be influenced by the rate of initial receptor-G protein coupling (i) or through modulating the G protein conformational changes that facilitate nucleotide exchange (ii). **b**, Labeling sites R90C (green) and E238C (magenta) shown on superimposed Gα_i_ structures in the inactive (GDP-bound) state (PDB: 1GP2) and in the receptor-coupled (nucleotide-free) state with an open AHD (PDB: 7L0Q). **c**, Ligand efficacy profiles determined from Gα_i_Δ6-90C/238C GTP turnover assay (Methods). Error bars denote 95% c.i., three technical replicates; *p < 0.023, **p < 0.004 and p > 0.058 (ns), two-sided, unpaired t-test with Bonferroni correction. BUP, buprenorphine; MOP, morphine; LFT, lofentanil. **d**, Schematic of smFRET experiments. **e**, Example single-molecule fluorescence (top) and FRET (bottom) time traces of surface-tethered μOR-G_i_ bound to full agonist lofentanil, displaying transitions between a low-FRET, open state and a high-FRET, closed state. **f**, Transition density plot displaying the mean FRET values before (x-axis) and after (y-axis) each transition for lofentanil-μOR-G_i_ (scale bar, 10^−3^ transitions per bin per second; *n*_*t*_total transitions).

The most significant conformational change observed for G protein activation by a GPCR is the separation of its Ras-like and α-helical domains (RLD and AHD) in the Gα subunit (Fig. 1a,b), as revealed by early double electron-electron resonance (DEER) spectroscopy^10^ and X-ray crystallography^11^ studies. In all inactive-state crystal structures, the AHD – a six α-helical bundle (αA-αF) – packs against the RLD, burying the bound GDP in the interface between the domains^12^. Following receptor coupling and GDP release, this interface opens by tens of angstroms^10,11,13–15^, resulting in an intrinsically dynamic AHD that has largely eluded high-resolution structural characterization^16^.

Multiple lines of evidence suggest that AHD movements are critical for facilitating GDP-GTP exchange across all G protein isoforms, yet their role in signaling efficacy remains poorly understood^17^. While molecular dynamics (MD) simulations indicate that domain separation alone may not be sufficient for nucleotide exchange, a variety of functional^18–20^, spectroscopic^13,21,22^, and computational^23,24^ studies demonstrate that AHD movements are tightly linked to both GDP release and the subsequent binding of GTP^16,25,26^. Here, we use total internal reflection fluorescence (TIRF) smFRET imaging to directly monitor AHD motions in G_i1_ proteins bound to μ-opioid receptor (μOR) activated by diverse orthosteric ligands, both in the absence and presence of nucleotide. Our results reveal how ligand-specific signals are allosterically encoded in the G protein conformational ensemble; a pattern that generalized to G_i_ bound to neurotensin type 1 receptor (NTSR1). These data suggest a mechanistic basis for opioid signaling efficacy that extends beyond simple receptor-G protein coupling efficiency.

## Results

Donor and acceptor fluorophores (LD555^27^ and JFX673^28^, respectively) were stochastically labeled on the AHD (R90C) and RLD (E238C) of a previously reported minimal cysteine Gα_i1_ mutant^10,29^ (Fig. 1b and Extended Data Fig. 1a; Methods). This construct, Gα_i_Δ6-90C/238C, exhibited a WT-like pattern of ligand efficacy-dependent GTP turnover with μOR in vitro (Fig. 1c), and LD555/JFX673-labeled Gα_i_Δ6-90C/238C heterotrimers efficiently coupled to agonist-bound μOR, forming stable, nucleotide-depleted ternary complexes (Extended Data Fig. 1b). For TIRF imaging (Methods), these μOR-G_i_ ternary complexes were specifically immobilized on quartz slides with an M1 fab fragment (Fig. 1d and Extended Data Fig. 1c), remaining functional on the surface as tested by the ability for GTPγS-induced G protein dissociation (Extended Data Fig. 1d). Time-dependent changes in AHD position were captured from smFRET trajectories of individual, immobilized μOR-G_i_ complexes, which displayed reversible transitions (on the order of 1 s^-1^) between low- and high-FRET states (Fig. 1e,f). These dynamics reflect spontaneous domain separation and re-engagement consistent with the various poses observed in previous negative stain and cryo-electron microscopy (cryo-EM) studies of a receptor-G protein complex^16,26^.

### Ligand-induced AHD opening

We examined the impacts of μOR bound to ligands with varying signaling efficacies (Fig. 1c), as well as the unliganded (apo) receptor, on AHD dynamics in coupled G_i_ heterotrimers (Fig. 2a and Extended Data Fig. 2a). These ligands included strong full agonists lofentanil and BU72^30^, the endogenous enkephalin-like full agonist DAMGO, the partial agonist morphine, and weaker partial agonists buprenorphine and mitragynine pseudoindoxyl (MP). Population FRET efficiencies for all ligands showed two predominant states, centered at ∼0.27 and ∼0.62, along with a minor ∼0.47 FRET population that remained constant across all samples. At μOR bound to full agonists, occupancy of the low-FRET (0.27) state increased relative to partial agonists, whereas that of the high-FRET (0.62) state decreased (Fig. 2a). This ligand-induced low-FRET occupancy was strongly correlated with in vitro GTP turnover activity (a direct measure of efficacy) (R^2^ = 0.97; Fig. 2b), indicative of an open Gα conformation like that seen in active-state complex structures^11,14^ (see Fig. 1b). Likewise, unliganded (apo) μOR was the least efficient at stabilizing this low-FRET, open conformation, in line with its slow GTP turnover under basal conditions (Fig. 2a,b). In addition, the apo complex exhibited a small population (∼10%) from excursions to >0.7 FRET, a state which the AHD rarely sampled under liganded conditions (Fig. 2a and Extended Data Fig. 2a).

**Fig 2.**
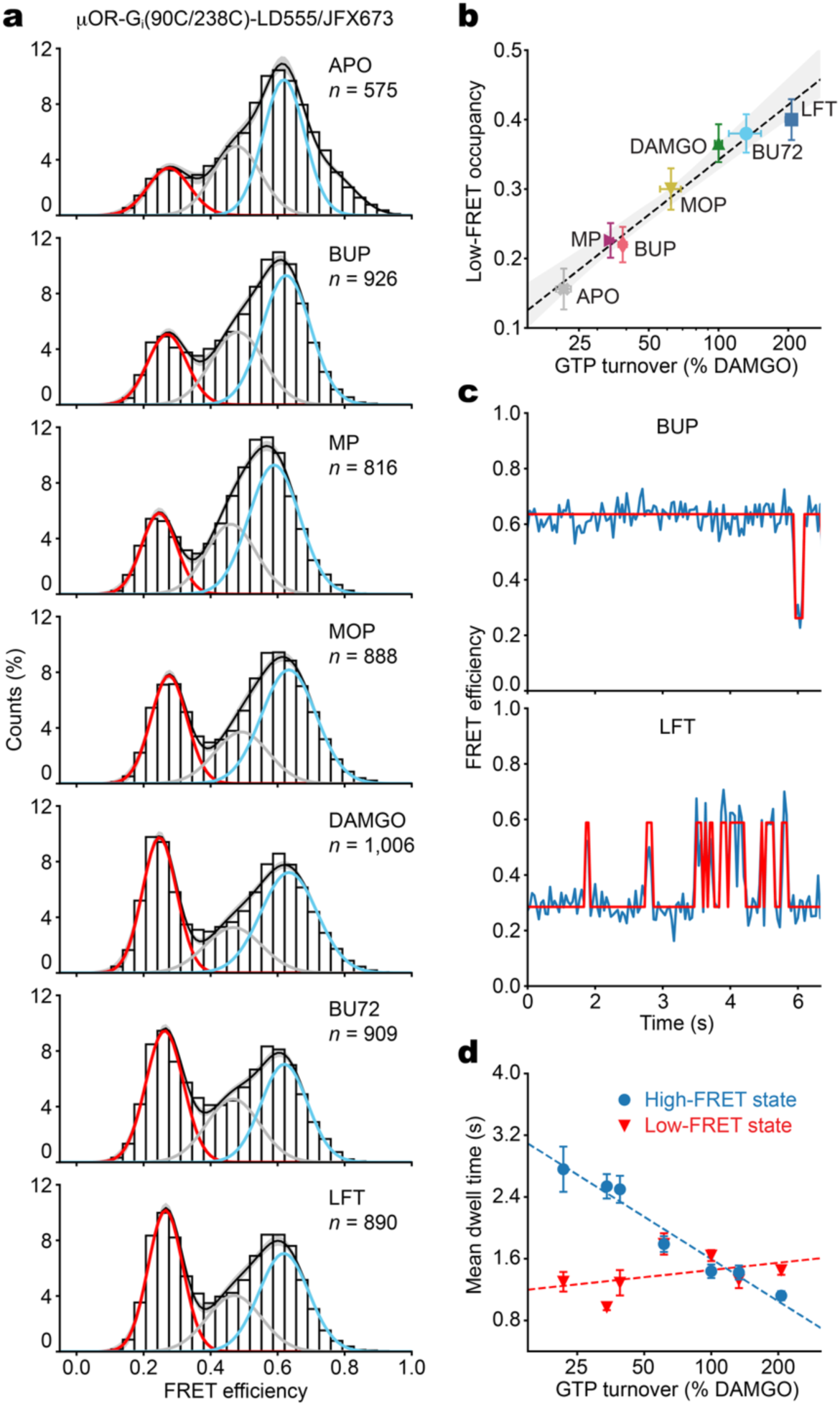
AHD opening is ligand efficacy-dependent. **a**, Population FRET efficiency histograms for surface-tethered μOR-G_i_ ternary complexes with three-state gaussian mixture model (GMM) fits showing the open (red) and closed (cyan) states (*n*, number of traces). Error bands, s.d. of GMM fits from 100 bootstrap samples of the FRET traces. **b**, Scatter plot of ensemble average low-FRET, open-state occupancies and mean GTP turnover activities observed for each ligand with log-linear fit (dashed line; R^2^ = 0.97). Errors represent 95% c.i. **c**, Example FRET (blue) and predicted state (red) traces for partial (top; buprenorphine) and full (bottom; lofentanil) agonists. **d**, Mean dwell times in the low-FRET, open and high-FRET, closed states; data is mean ± s.d., 100 bootstrap samples of the idealized traces.

To investigate the kinetics of AHD motions within μOR-G_i_, we used hidden Markov modeling (HHM) to idealize the traces and quantify transitions between high-and low-FRET states (Figs. 1e and 2c; Methods). The occupancy of the mid-FRET (∼0.47) state was not ligand-dependent (Fig. 2a) and was sampled transiently, with nearly half of its occurrences lasting two frames or less (≤ 100 ms). Hence, the mid-FRET state may reflect an unstable conformational intermediate and/or result from time-averaging of rapid (> 10 s^-1^) low-to-high FRET excursions at the current imaging speed. Because neither mid-FRET transitions nor distributions could be clearly distinguished from the high-FRET state at the ensemble level (Extended Data Fig. 2b and Fig. 2a), we analyzed the kinetics as a two-state system, where the mid- and the high-FRET states are collapsed into a single, broadly distributed (∼0.4-0.6) closed conformation.

This analysis revealed that increasing ligand efficacy at μOR enhances the overall level of dynamics within coupled G_i_ heterotrimers (Fig. 2c and Extended Data Fig. 2b,c), primarily by reducing the duration that the AHD dwelled in the closed, high-FRET state (Extended Data Fig. 2c). Mean dwell times in this high-FRET state (*τ*_high_) depended strongly on ligand efficacy (R^2^ = 0.96; Fig. 2d), with the apparent rate of AHD opening being approximately 2.5-fold faster for lofentanil-compared to apo-occupied complexes (ca. 0.9 s^-1^ and 0.36 s^-1^, respectively). Conversely, dwell times in the low-FRET state were similar (ca. 1.4 s on average) across all ligands tested (R^2^ = 0.17; Fig. 2d and Extended Data Fig. 2c). Thus, ligand-dependent shifts in the AHD’s conformational equilibrium (Fig. 2a) stem from the destabilization of its closed state, rather than stabilization of the open, domain-separated conformation. This indicates that efficacy is, at least in part, dependent on the ability of the ligand to allosterically initiate domain separation rather than merely to promote receptor conformations that affect initial G protein coupling.

While domain separation is key for receptor-catalyzed GDP release, the observed high-to-low FRET transition rates are slower (approximately 10-fold) than expected from G protein activation measurements in living cells (< 100-300 ms)^31,32^. However, the absolute values of these rates may be influenced by experimental conditions, such as the detergent micelle, cysteine mutations, temperature, and/or limitations in imaging speed and fluorescent probes. Parallel smFRET imaging of NTSR1-G_i_ ternary complexes (Methods), which couple promiscuously in vitro, yielded a similar correlation between agonist-induced low-FRET occupancy and GTP turnover (R^2^ = 0.89; Extended Data Figs. 3 and 4; cf. Fig. 2b). The addition of SBI-553, an allosteric modulator shown to elicit an effector-dependent response^33^, shifted the AHD toward the open state proportionally to its positive modulation of GTP turnover in vitro (Extended Data Fig. 3b). These findings suggest that AHD displacement may be a general determinant of ligand efficacy across G protein-receptor pairs, consistent with previous bioluminescence resonance energy transfer (BRET) studies showing distinct G protein conformational signatures for different calcitonin and α2-adrenergic receptor ligands^22,34^.

### G protein dynamics in the basal state

In basal (inactive)-state structures of heterotrimeric G proteins, the Gα AHD tightly encloses the bound GDP^35,36^. Yet spontaneous domain separation must occur to permit GDP release, as G proteins still slowly exchange nucleotide even without GPCR catalysis^37^. To investigate these basal dynamics, we immobilized labeled G_i_ heterotrimers via an N-terminal histidine-tag and imaged their AHD motions in a physiological concentration of GDP (30 μM^38^) (Extended Data Fig. 5a; Methods). Compared to receptor-bound G protein, we observed more conformational heterogeneity (≥ 3-4 distinct states) and reduced average interdomain separation, primarily due to the AHD sampling an additional (inactive, high-FRET) state centered at ∼0.77 (Extended Data Fig. 5b-d). This inactive state corresponds to an interdye distance consistent with full domain closure (Δopen-closed ≈ 28 Å; Methods) and likely resembles the GDP-bound conformation seen in the inactive-state G_i_ crystal structure^35^ (see Fig. 1b). These data predict that even in its “closed,” 0.62 high-FRET state, the receptor coupled G_i_ AHD remains partially displaced (ca. 5-8 Å). Elimination of the fully-closed (0.77 FRET) conformation upon agonist-stimulated receptor-G protein complex assembly (Fig. 2a and Extended Data Fig. 3a) is likely related to the mechanism of GDP release (or simply the absence of bound GDP).

Under basal conditions, individual G_i_ AHD exhibited reduced dynamics but still transitioned to the open state (ca. 0.15 s^-1^; Extended Data Fig. 5c,d), consistent with prior MD simulations^23^. These transitions predominantly traveled through a mid-FRET (∼0.5) intermediate state (Extended Data Fig. 5d), suggesting that a partially open AHD may mediate transitions between GDP-bound and GDP-unbound conformations. Introducing an alanine substitution at residue R144 in the α subunit (Extended Data Fig. 5e), previously shown to destabilize the RLD-AHD interaction and enhanced basal nucleotide exchange^37,39^, shifted the AHD from the fully closed to open states, resulting in dynamics similar to, though still slower than, the receptor-bound G protein (Extended Data Fig. 5f-h).

### Nucleotide effects on intact μOR-G_i_ complexes

As described above, the AHD can adopt at least four distinct FRET states: (i) a low-FRET, open state (∼0.27) that correlates with ligand efficacy; (ii) a short-lived, intermediate state (∼0.47); (iii) a high-FRET, closed state (∼0.62) while the G protein is receptor-bound; and (iv) an inactive, GDP-bound closed state (∼0.77) seen basally in uncoupled G_i_. The FRET states observed in receptor-coupled G proteins (0.27, 0.47, and 0.62) could not be directly linked to stable nucleotide binding, as GTP or GDP delivery caused rapid dissociation from the receptor, precluding equilibrium observation of bound states (Fig. 3a). However, GMP and pyrophosphate (PPi) (Extended Data Fig. 6a), which together mimic GTP binding in the nucleotide pocket of small G proteins^40^, did not significantly dissociate the complex (Fig. 3a,b).

**Fig 3.**
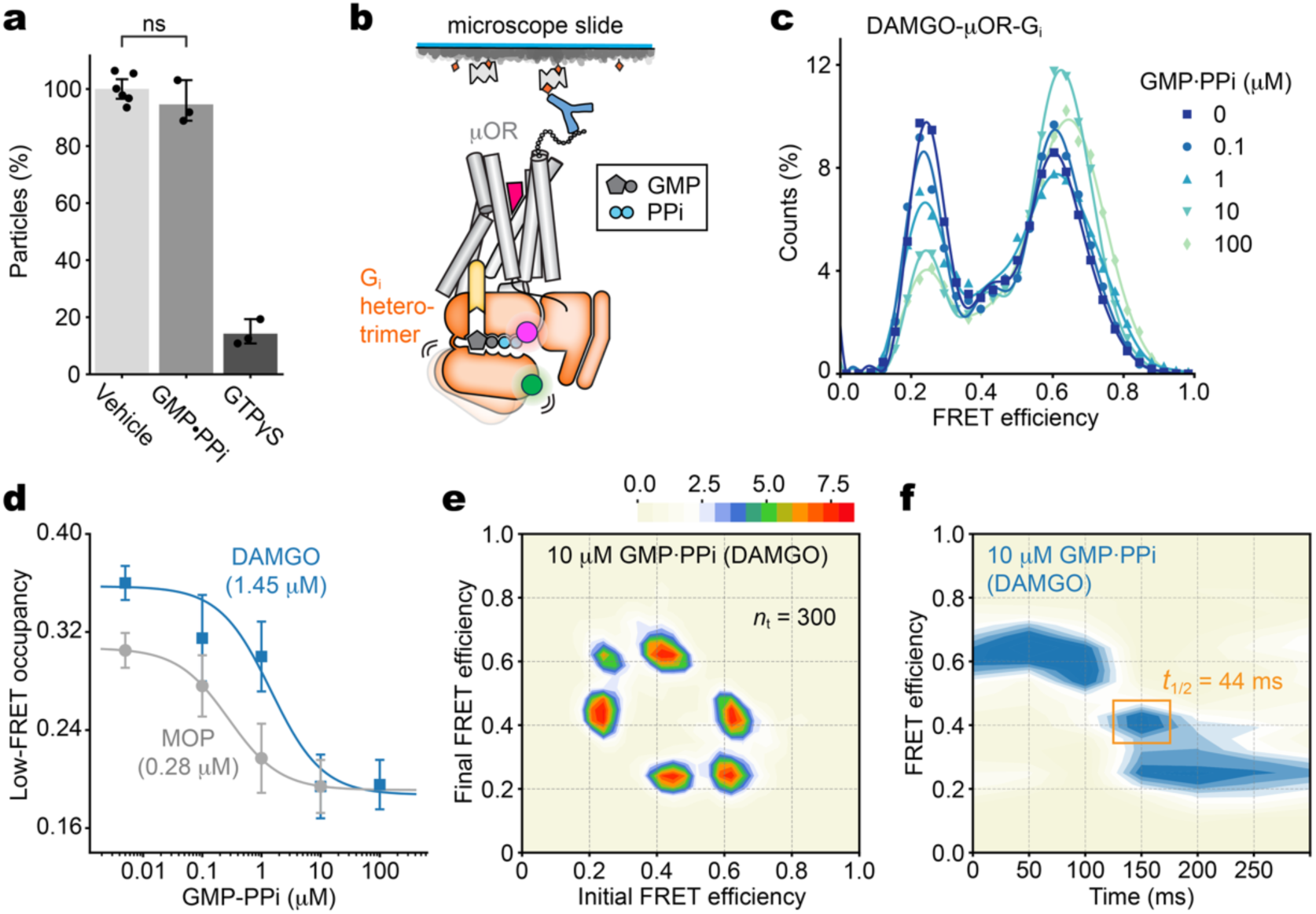
Nucleotide binding promotes AHD closure through an intermediate. **a**, GTPγS, but GMP with PPi, dissociates G protein from surface-tethered μOR. Error bars, mean ± 95% c.i. (n = 3-6 movies of the same sample) of FRET particles remaining after delivery of 10 μM nucleotide or buffer (vehicle); p = 0.23 (ns), two-sided, unpaired t-test. **b**, Schematic of single-molecule GMP·PPi titration experiments. **c**, Population FRET efficiency histograms (symbols) with b-spline fits (lines) for DAMGO-bound μOR-G_i_ in the presence of increasing concentrations of GMP·PPi. **d**, Apparent EC_50_ values measured from GMP·PPi-induced ensemble average low-FRET occupancy for DAMGO (blue) and morphine (gray); error bars, s.d. of 100 bootstrap samples of the FRET traces (3,341 molecules total across all conditions). Lines are fits to dose response functions (Hill slope = 1.0). **e,f**, Transition density plot (**e**) for DAMGO with 10 μM GMP·PPi (scale bar, 10^−3^ transitions per bin per second; *n*_*t*_total transitions) and contour plot of synchronized high-to-low transitions (**f**) showing a short-lived dwell in a mid-FRET state (yellow box, width ≈ one frame) (cf. Extended Data Fig. 6e).

**Fig 4.**
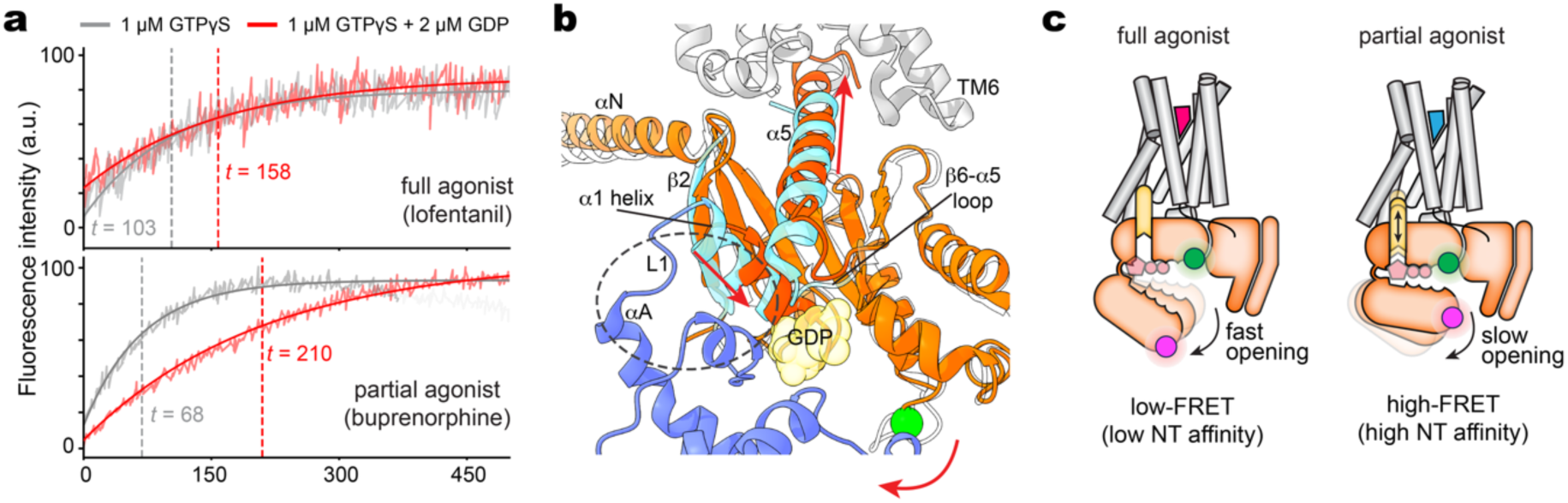
Model for the role of G protein dynamics in ligand-dependent signaling. **a**, Binding of BODIPY-conjugated GTPγS (GTPγS-BDP-FL) to nucleotide-free μOR-G_i_ containing unlabeled Gα_i_Δ6-90/238C subunit was monitored via fluorescence dequenching for lofentanil (top) and buprenorphine (bottom). Normalized kinetic spectra in the absence (gray) or presence (red) of GDP were fit using exponential association to obtain the mean association time (*t*; dashed lines). **b**, Engagement of μOR intracellular surface by the Gα_i_ α5 helix destabilizes the α1 helix and disrupts the interdomain interface (dashed outline) composed of αF, α1, L1, and the n-terminal segment of αA. Structures shown: 1GP2 (blue) and 7T2H (orange). **c**, Schematic of main FRET states and their proposed relation to α5 insertion efficiency and nucleotide (NT) binding. Full agonists favor a more open G protein complex (left), lowering nucleotide affinity. Partial agonists promote greater domain closure (right), increasing the energetic barrier for nucleotide release and reducing the likelihood of progressing to the nucleotide-free complex required for activation.

Using this approach with surface-tethered μOR-Gi complexes activated by DAMGO (Fig. 3c-f and Extended Data Fig. 6b-e), we observed a concentration-dependent shift from open to closed states induced by 0.1-100 μM GMP·PPi (Fig. 3c and Extended Data Fig. 6b). This is consistent with previous reports that domain closure is required for triphosphate nucleotide binding^25,41^ and reflects GMP·PPi binding through stabilization of the high-FRET closed conformation rather than an acceleration of domain closure (Extended Data Fig. 6c). The GMP·PPi-induced transitions also exhibited ligand-specific differences: complexes with the lower-efficacy ligand morphine (Fig. 3d and Extended Data Fig. 6f-h) showed heightened sensitivity to GMP·PPi (EC_50_ = 0.28 μM) compared to those with DAMGO (EC_50_ = 1.45 μM) (Fig. 3d). This difference aligns with the ligand-dependent energetic biases for AHD closure, as evident from their conformational equilibriums before nucleotide binding (Fig. 2a). Importantly, even at the highest GMP·PPi concentrations tested (≥10 μM), transitions to the basal (∼0.77 FRET) state remained rare (Extended Data Figs. 6b,f and 7a,b), suggesting receptor coupling restricts access to the fully closed conformation of the AHD regardless of nucleotide.

Further analysis of individual smFRET traces (Extended Data Figs. 6d,g and 7a,b) revealed stepwise transitions through a partially open, mid-FRET state (∼0.4–0.5) within both DAMGO- and morphine-bound complexes (Fig. 3e and Extended Data Fig. 6h). GMP·PPi-induced stabilization of this intermediate was weaker in DAMGO-bound complexes, which unlike morphine, maintained some apparent direct transitions between low- and high-FRET states even at saturating nucleotide concentrations (Fig. 3e). However, synchronization analysis revealed a brief (1-2 frame) mid-FRET dwell during many of these domain separation events (Fig. 3f, cf. Extended Data Fig. 6e). We had also observed stepwise transitions in heterotrimeric G_i_ with GDP present (Fig. 5d), consistent with a prior smFRET study linking the partial domain separation in the mid-FRET state to an intermediate level of nucleotide affinity^21^.

Together, these findings suggest a mechanism where nucleotide contacts are broken or formed progressively rather than all at once, providing a free-energy-efficient pathway for nucleotide exchange via a transient intermediate step^42,43^. Similar accessibility of this binding-competent intermediate in the nucleotide-free state seen across ligands (Fig. 2a) may allow for selective modulation of GDP release (i.e., by tuning high-FRET state stability) without as drastically affecting GTP binding between ligands.

## Discussion

We have shown that opioid ligand efficacy extends beyond initial receptor-G protein coupling to include the modulation of G protein conformational equilibria within the fully formed signaling complex. These results demonstrate that the G protein conformation can be dynamically tuned by the GPCR, sampling a continuum of distinct states rather than merely discrete ‘off’ or ‘on’ conformations (Fig. 2b). While these insights advance our understanding of opioid drug action – informing efforts to develop safer analgesics^44–47^ – they also suggest a broader framework for how interplay between receptor and effector conformational states can influence the signaling response amplitude, which has clinical implications across GPCR targets^48,49^.

Our findings reveal that ligand-dependent conformational changes propagate from the receptor to the engaged G protein, modulating its AHD between open, closed, and transient intermediate states. Consistent with the loss of contacts upon separation of the Gα domains, measurements showed that the high-FRET (closed) state corresponds to higher nucleotide affinity, while the low-FRET (open) state possess reduced affinity (Fig. 3c,d). This aligns with prior findings that mutations to the interdomain interface or hinge regions accelerate basal nucleotide exchange^37,50–52^, and that increased interdomain motion correlates with G protein activation^53–55^. Thus, the energetic bias towards AHD opening observed in full agonist-bound complexes (Fig. 2a and Extended Data Fig. 3a) suggests a reduction in nucleotide affinity, driven by the shift in the G protein conformational equilibria.

To test this, we measured GTPγS binding to nucleotide-free μOR-G_i_ complexes by fluorescence dequenching (Methods). Buprenorphine-occupied complexes exhibited 1.5-fold faster GTPγS binding than those bound to the full agonist lofentanil (Fig. 5a), consistent with extensive earlier evidence that higher-efficacy ligands reduce the nucleotide affinity of coupled G proteins^5,56–61^. Although this initially appears to contradict in vivo signaling patterns, measurements under more physiological conditions – where both GDP and GTP are present – showed a reversal in binding kinetics (Fig. 5a), with GDP competition dramatically slowing GTPγS binding to buprenorphine-bound complexes^62^.

In the cytoplasm, where GTP is ∼10-fold more abundant than GDP, G protein activation is primarily limited by GDP release, as GTP loading occurs rapidly afterward. Full agonists promote AHD opening, facilitating GDP release and efficient GTP binding. In contrast, partial agonists restrict open-state occupancy, favoring GDP retention or rebinding, which prolongs its residence time in the interdomain pocket. This competitively inhibits GTP binding and thereby delays G protein activation, ultimately explaining why partial agonists exhibit lower signaling efficacy despite faster intrinsic GTPγS binding rates (Fig. 5a). The relative difference between ligands is also greater for GDP than for GTP binding, further suggesting that G protein dynamics are primarily selected to regulate the GDP release step.

At a structural level, these ligand-induced G protein conformational changes require allosteric communication spanning over 70 Å from the receptor’s orthosteric site to the G protein AHD. Differences in G protein dynamics observed with different orthosteric ligands can be understood as alterations in the complementarity between the binding surfaces where the RLD and AHD meet. While GPCR-mediated G protein activation is known to involve remodeling of the α5 helix in the Gα subunit^63–66^, studies also link this helix to AHD opening^10,23,67^ – minimally through the α1 helix in the RLD^68,69^ (Fig. 5b). The α1 helix directly connects to the AHD via the αA helix and helps stabilizes the inactive G protein conformation through conserved interdomain contacts with αF and the hinge region^70^. During GPCR engagement, α5 moves away from the nucleotide-binding pocket toward the intracellular vestibule of the receptor, causing rotation and partial disorder of α1. These rearrangements sterically disrupt the interdomain interface, opening the AHD and priming the ternary complex for GDP release^13,15,66^ (Fig. 5b).

Recent smFRET and DEER studies of β2 adrenergic and μOR receptors showed that full agonists promote greater outward movement of transmembrane helix 6 (TM6)^5,6^, expanding the intracellular vestibule where α5 binds. This TM6 displacement, along with other ligand-dependent changes to the receptor’s cytosolic surface, may facilitate AHD opening by making it more energetically favorable for G protein to bind^15,71–73^. In contrast, partial agonists have been shown to induce less efficient α5 insertion^74^, which could weaken α1-mediated disruption of the Gα interdomain interface and prolong the dwell time in the high-FRET, high-nucleotide-affinity state preceding GDP release (Fig. 5c). This mechanism helps explain an earlier observation that partial agonists at β2-G_s_ act to stabilize transient, GDP-bound GPCR-G protein intermediates^5^, which could function as kinetic traps retarding productive complex assembly and activation. As for the low-affinity open state, the similar low-FRET dwell times observed across opioid agonists (Fig. 2d) suggest that once the AHD is fully open, ligand identity has minimal influence on its dynamics – possibly because the AHD is affixed against Gβ, as seen in a recent cryo-EM structure^14^.

Unlike real-time insights from smFRET, resolution of the AHD in cryo-EM studies of GPCR-G protein complexes is limited by the conformational heterogeneity imaged upon freezing^16,55^. As such, it may be unsurprising that structures with ligands of varying efficacies tend to show minimal differences in Gα^57,75,76^, capturing only lowest-energy conformations while even subtle changes at the interdomain interface can significantly alter domain separation dynamics^70^. By revealing that GPCRs allosterically control nucleotide exchange through biasing the conformational equilibria of the AHD, our work provides mechanistic evidence for how opioid and other GPCRs function as guanine nucleotide exchange factors^17^. This dynamic framework provides a more complete picture of GPCR signal transduction and suggests new strategies for developing opioid therapeutics with optimized efficacy profiles and potentially fewer adverse side effects.

## Methods

### Minimal cysteine Gα_i_ expression, purification, and labeling

Mutations R90C and E238C were introduced into a previously published^29^ minimal cysteine background (C3S, C66A, C214S, C350S, C325A, C351I) of human Gα_i1_ containing an N-terminal 6x histidine-tag prior to a rhinovirus 3C protease site. This construct (Gα_i1_Δ6-90C/238C) was expressed in BL21 cells using pET21a and purified as previously described^77^. Briefly, cells grown in Terrific Broth were induced with 0.5 mM IPTG at an OD_600_ of 0.6 and harvested after 15-20 hours at room temperature. The harvested cells were resuspended in lysis buffer (50 mM HEPES pH 7.4, 100 mM sodium chloride, 1 mM magnesium chloride, 50 μM GDP, 5 mM beta-mercaptoethanol, 5 mM imidazole, lysozyme, and PIs) and disrupted via sonification. Intact cells and cell debris were removed by centrifugation and Gα_i_ was purified from the supernatant using Ni-NTA resin. The protein was eluted using 200 mM imidazole in lysis buffer and dialyzed with 1:1000 w/w 3C protease at 4 °C overnight to cleave the histidine tag. 3C protease, cleaved histidine tag and unreacted protein was removed using fresh Ni-NTA and the flow-through containing purified Gα was concentrated and polished using size exclusion chromatography (SEC) on a Superdex 200 10/300 GL column (Cytiva) in G protein buffer (20 mM HEPES pH 7.4, 100 mM sodium chloride (NaCl), 1 mM magnesium chloride, 20 μM GDP, and 100 uM TCEP). For preparation of His-tagged Gα_i_, the Ni-NTA elute was concentrated and further purified using SEC without the intervening 3C cleavage step. The R144A mutant of Gα_i1_Δ6-90C/238C was purified with an intact His-tag in the same manner. Aliquots of Gα_i_ were flash frozen in liquid nitrogen at a concentration of approximately 200 μM in 20 % glycerol as a cryoprotectant.

For site-specific labeling, fluorophores were attached stochastically to C90 and C238 in Gα_i_Δ6-90C/238C under conditions designed to minimize off-site labeling (Extended Data Fig. 1a). Aliquots of frozen, purified Gα were brought to room temperature and diluted to a concentration of 30 μM in G protein buffer containing 50 μM GDP. Maleimide-conjugated LD555 (Lumidyne Technologies) and JFX673peg_4_ (Janelia Fluor) were added simultaneously, at a 2- and 1.7-fold molar excess, and incubated in the dark at room temperature for 30 minutes. The reaction was quenched by 5 mM L-Cysteine (Sigma) before labeled protein was purified from the unreacted dyes using a Superdex 200 column in buffer containing 20 mM HEPES pH 7.4, 100 mM NaCl, 1 mM MgCl_2_, 20 μM GDP, and 100 uM TCEP. On average, labeling efficiency was >80%, with an LD555:JFX673 ratio of <1. SEC-pure, dye-labeled Gα_i_ was concentrated, and flash frozen at approximately 60 μM for future use.

### Purification of Gβγ

Gβγ heterodimer was expressed in *Tni* cells using a baculovirus generated by the BestBac (Expression systems) method and purified as described previously^23^. Briefly, we used a single virus containing the genes for both the Gβ_1_ and Gγ_2_ subunits. A N-terminal 6x histidine tag followed by a 3C protease site was contained in the sequence of Gβ to be used for Ni affinity purification. Gβγ heterodimer was extracted from the membrane using a dounce homogenizer in solubilization buffer containing 20 mM HEPES pH 7.4, 100 mM NaCl, 1.0% sodium cholate, 0.05% dodecylmaltoside (DDM, Anatrace), 5 mM beta-mercaptoethanol, and PIs. After removing insoluble debris, the protein was washed with a solubilization buffer in batch and gradually exchanged from sodium cholate to 0.1% DDM on Ni-NTA resin. The elute was dialyzed with 3C protease to cleave the histidine tag as we described for Gα_i_. Cleaved Gβγ was dephosphorylated and further purified on a MonoQ 10/100 GL column (GE Healthcare) using a linear gradient of NaCl (50-250 mM). The main peak containing prenylated Gβγ was dialyzed into 20 mM HEPES pH 7.4, 100 mM NaCl, and 0.02% DDM and flash frozen in liquid nitrogen at a concentration of 150 μM with 20% glycerol as a cryoprotectant.

### Formation dye-labeled G_i_ heterotrimer

Purified Gβγ heterodimer was exchanged into buffer containing 20 mM HEPES pH 7.4, 100 mM NaCl, 1 mM magnesium chloride, 50 μM GDP, and 100 uM TCEP using a 30k MWCO centrifugal protein concentrator (Thermo Scientific) and mixed with approximately 0.25 molar equivalents of dye-labeled Gα_i_Δ6-90C/238C-6x histidine (2.5 μM Gαi final concentration). Buffer exchange was performed until the final concentration of DDM during heterotrimer formation was << the sub critical micellar concentration (CMC). After incubation at room temperature for 0.5 hours, unbound Gβγ heterodimers were removed coincident with surface immobilization for smFRET imaging by pulling down the heterotrimer on microscope slides coated in anti-His epitope tag antibodies. Unlabeled, wild-type human G_i_ heterotrimer (Gα_i1_β_1_γ_2_) was co-expressed and purified from *Hi5* insect cells as previously described^23^.

### Purification of μOR

Mouse μOR was expressed in *sf9* insect cells using the baculovirus method (Expression systems) in the presence of the opioid antagonist naloxone (Sigma Aldrich). We used a modified *Mus musculus* sequence containing an N-terminal Flag tag and an 8x histidine tag in front of a rhinovirus 3C protease site on the C-terminus. μOR was expressed and purified in the presence of 10 μM naloxone as has been previously described^78^. Briefly, *sf9* cells were infected at a density of 4 × 10^6^ ml^-1^ and incubated for 48 hours at 27 °C. Cell membrane was solubilized in DMM and 3-[(3-cholamidopropyl)-dimethylammonio]-1-propane sulfonate] (CHAPS, Anatrace) and purified by immobilized metal affinity chromatography using Ni-NTA resin. The elute was loaded onto M1 anti-Flag immunoaffinity resin and gradually exchanged into a buffer containing naloxone, 0.01% lauryl maltose neopentyl glycol (L-MNG, Anatrace) and 0.001% cholesterol hemisuccinate (CHS, Steraloids). The receptor was further purified on SEC in a buffer containing no ligand, 25 mM HEPES pH 7.4, 100 mM NaCl, 0.02% L-MNG and 0.002% CHS. The peak containing the now unliganded μOR (Extended Data Fig. 1b, dashed line) was concentrated to ∼100 μM, supplemented with 20% glycerol, and flash frozen for later use.

### Assembly of μOR-G_i_

To prepare μOR-G_i_ ternary complexes for smFRET, μOR was equilibrated in a 5-10 molar excess of opioid agonist for 10 minutes at room temperature. After pre-incubation of untagged Gα_i_ and Gβγ on ice for 30 minutes, they were mixed with liganded receptor at a 1:1.2:1 (Gα_i_:Gβγ:μOR) molar ratio. The final concentration of receptor during complexing was ∼2.5 μM and the buffer adjusted to contain 20 mM HEPES pH 7.4, 100 mM NaCl, 0.01% L-MNG, 0.001% CHS, 100 μM TCEP, 50 μM GDP and 1 mM MgCl_2_. Complex assembly progressed for one hour at room temperature to reach steady state, after which 25 mU ml^-1^ Apyrase (New England Biolabs) was added and incubated overnight at 4 °C. The apyrase-stabilized complex was supplemented with 2 mM CaCl_2_ and loaded onto a M1 anti-Flag immunoaffinity column at room temperature. The column was washed with 20 column volumes (CV) of complex buffer (20 mM HEPES pH 7.4, 100 mM NaCl, 0.01% L-MNG, 0.001% CHS and 1-10 μM ligand) containing 2 mM CaCl_2_ before the protein was eluted using 200 μg ml^-1^ Flag peptide (DYKDDDDK) and 2 mM EDTA. The elute was further purified using a room temperature Superdex 200 10/300 SEC column in complex buffer containing 0.1-1 μM ligand. Peak fractions corresponding to monodispersed μOR-G_i_ were concentrated to ∼3 μM and supplemented with 20% glycerol before flash freezing in liquid nitrogen. Single-use aliquots were stored at -80 °C until imaging. Nearly identical FRET efficiency distributions were obtained using complexes that had not been frozen (data not shown). Given that the unliganded μOR-G_i_ complex showed reduced stability at room temperature, all chromatography and sample preparation for the apo was performed at 4 °C.

### Purification and assembly of labeled NTSR1-G_i_

Full-length human NTSR1 was expressed and purified as previously described^79^. In brief, an NTSR1 sequence modified with an N-terminal Flag tag followed by an octa-histidine tag was expressed in *sf9* cells using a baculovirus. Solubilized cells were purified using sequential Ni-NTA, anti-Flag and SEC columns into a final buffer containing 20 mM HEPES, pH 7.4, 100 mM NaCl, 0.0025% L-MNG, 0.00025% CHS and 0.1 μM NTS_8–13_ (acetate salt, Sigma). Peak fractions were concentrated to 200 μM and flash frozen and stored at -80 °C.

Ternary complexes of NTSR1-G_i_ were assembled and purified as described for μOR with a few modifications. Purified NTSR1 in 0.1 μM NTS_8–13_ was loaded onto an anti-Flag column and exchanged into different orthosteric ligands (NTR1-17 or CGX-1160) by washing for 0.5 hours (∼15 CV) at room temperature with buffer containing 1 μM of the new ligand and 20 mM HEPES, pH 7.4, 100 mM NaCl, 0.0025% L-MNG, and 0.00025% CHS. Flag peptide and EDTA were removed from the elute using a 7k MWCO desalting spin column (Thermo Scientific) before the NTSR1 was concentrated and used for complexing. To couple to the receptor, labeled Gα_i_Δ6-90/238C and Gβγ were first assembled into heterotrimer as described above and were mixed with NTSR1 in an ∼1:1 ratio following incubation. Complexing then progressed as for liganded μOR-G_i_ using buffers containing NTSR1 ligands.

### GTP turnover assays

Analysis of GTP turnover was performed using a modified GTPase-Glo™ assay (Promega) as described previously^5^. We started the reaction by mixing purified, unliganded or ligand-bound μOR or NTSR1 and G_i_ protein in an assay buffer containing 20 mM HEPES, pH 7.5, 100 mM NaCl, 0.01% L-MNG, 0.001% CHS, 100 µM TCEP, 10 mM MgCl2, 10 µM GDP, 5 µM GTP and saturating concentrations of receptor ligands. After incubation for 90 minutes, reconstituted GTPase-Glo™ reagent was added to the sample and incubated for 30 minutes at room temperature. Luminescence was measured after the addition of detection reagent and incubation for 10 minutes at room temperature using a SpectraMax Paradigm plate reader. Measurements were performed in triplicate on the same plate and reported as the mean relative light units (RLU) normalized to DAMGO or NTS_8-13_. Ligand efficacy profiles were determined from the GTP turnover activity stimulated by μOR or NTSR1, using Δ6-90C/238C and WT Gα_i_ constructs, respectively. Basal G_i_-dependent GTP turnover was measured using buffer without any receptor and a 120-minute incubation period.

### smFRET measurements

Data was acquired using a lab-built prism-based TIRF microscope. LD555 (donor) and JFX673-peg_4_ (acceptor) labeled Gα_i_-Δ6-90/238C was excited using a 532 nm solid-state laser (Coherent OBIS™ LS Laser) set to 40 mW at the laser. Photons emitted from LD555 and JFX673 were collected through a 1.27 NA 60x water-immersion objective (Nikon) and split onto two cooled EMCCD cameras (Andor iXon Ultra 888) connected by a 640 nm dichroic mirror (ZT640rdc, Chroma) housed in a TwinCam™ (Cairn). All imaging was performed at 19 °C and movies were recorded with Andor Solis (i) software (Andor) at a 50 ms time resolution (20 frames s^-1^ (fps)). Microfluidic chambers for smFRET imaging were constructed from passivated quartz slides and coverslips using a modified protocol from Ha et. al^80^. Briefly, aminated glass surfaces passivated with mPEG-SVA (MW 5000; Laysan Bio Inc.) doped with 0.6% w/w biotin-PEG-SVA (MW 5000; Laysan Bio Inc.) were exposed to two additional rounds of passivation reactions using only mPEG-SVA.

For experiments with receptor-G_i_ ternary complexes, passivated imaging chambers were incubated for 5 minutes with 0.2 mg ml^-1^ NeutrAvidin (Thermo Scientific), followed by 5 minutes with 25 nM biotinylated anti-Flag M1 Fab fragment. Unbound M1 Fab and NeutrAvidin were washed out using imaging buffer (20 mM HEPES, pH 7.5, 100 mM NaCl, 0.01% L-MNG, 0.001% CHS, 2 mM CaCl_2_). Fluorophore-labeled μOR-G_i_ and NTSR1-G_i_ complexes were filtered using a 0.22 μm PVDF membrane (Millipore Sigma) and loaded onto the microscope slide at 200 pM by pulling down the N-terminal Flag tag on the receptor. Unbound complexes were washed out at an immobilization density of ∼0.1 molecules μm^2^. All experiments were performed in imaging buffer containing a saturating concentration of the orthosteric and/or allosteric ligand. For smFRET of basal G_i_ heterotrimer, NeutrAvidin-coated chambers were incubated for 5 minutes with 0.02 mg ml^-1^ biotinylated anti-mouse IgG1 antibody (Abcam) followed by 5 minutes with 0.02 mg ml^-1^ mouse anti-hexa-histidine epitope tag antibody (Abcam). Purified, LD555/JFX673peg_4_-labeled G_i_ with an uncleaved N-terminal hexa-histidine tag on the α subunit was loaded onto the microscope slide at ∼200 pM and measured in imaging buffer containing 30 μM GDP, 1 mM MgCl_2_ and 100 μM TCEP. Anti-his tag immobilization has been reported previously^81,82^ and was specific (data not shown).

### smFRET data analysis

Fluorescence time trajectories of single molecules were extracted from the movies and processed using a custom software suite implemented in MATLAB which is available on GitHub. Donor traces were scaled by a constant factor (γ) to account for the apparent unequal fluorophore brightnesses, calculated as γ = Δ*I*_A_/Δ*I*_D_, where Δ*I*_A_and Δ*I*_D_are the changes in donor and acceptor intensity after acceptor photobleaching, respectively^83^. FRET trajectories were calculated as (1 + (γ*I*_D_/*I*_A_))^-1^, where *I*_D_and *I*_A_are the background-corrected donor and acceptor fluorescence intensities at each frame. FRET traces lasting at least 0.5 s were selected for analysis that exhibited: (i) single-step photobleaching, (ii) overall anti-correlated donor and acceptor intensities, (iii) a signal-to-noise ratio (SNR) (mean/s.d. of the total fluorescence intensity before photobleaching) greater than 2, and (iv) acceptor photobleaching and/or low-FRET states with an average FRET of at least ∼0.2 (i.e., 1-2 s.d. from the center of the crosstalk distribution created by donor leakage into the acceptor channel). All traces were manually inspected for the final datasets, which had a geometric average SNR of ∼11 across all samples. Population histograms summed over the first 3 s of the selected FRET traces were fit to a mixture of 3-4 gaussian functions to derive the mean FRET values and occupancies of each state. Initial guesses for the states were obtained from fitting the full agonists. Errors for the gaussian mixture and state occupancies were calculated as the s.d. of 100 bootstrap samples of the FRET traces (95% c.i. were obtained from 1.96 multiplied by s.d.).

FRET traces were idealized using a two-or three-state hidden Markov model (HMM) as implemented in the vbFRET software^84^. The idealized FRET traces (viterbi) were clustered using a gaussian mixture to remove overfitting of fluctuations within the same FRET state, and the resulting state sequences were used to calculate transition density plots (TDPs) from transitions corresponding to ΔFRET_ideal_ > 0.1. The number of states *k* for each condition was determined based on the lowest value for which a *k* + 1 state HMM fit did not change the clusters in a TDP of the uncorrected traces. To explore the exchange kinetics between the fully open and closed FRET states within μOR-G_i_, the state sequences resulting from a two-state HMM and gaussian mixture fit were concatenated (“stitched”) and the dwell times in each state were extracted from the change points. Population dwell time distributions were calculated using a kernel density estimate (smoothing bandwidth = 2x Scott’s rule^85^) which was normalized to reflect the frequency of dwells in time bins along the x-axis^86^. The mean lifetime in each state was estimated as the integral of the normalized dwell-time survival curves restricted to < 30 seconds. Dwell times and their distributions are reported as the mean ± s.d. from 100 bootstrap samples of the FRET traces, where the resampling probability of each trace is weighted by its relative duration, as described previously^87^. As each bootstrapped sample has a different randomly selected order of traces, this source of variability is averaged over and reflected in the reported errors.

Synchronization plots of the high-to-low transitions in DAMGO-μOR-G_i_ were calculated from clustering the viterbi to two-states and selecting transitions from above to below the intermediate state center (∼0.45). While kinetics were analyzed with only γ correction, conversions of FRET efficiency to relative inter-dye distances utilized additional corrections for channel crosstalk and direct acceptor excitation, as outlined by Hellenkamp et al^88^. Figure preparation and downstream analysis of the selected FRET and idealized trajectories were performed using custom scripts in python.

### Bodipy-GTPγS binding assay

Binding of Bodipy conjugated GTPγS (GTPγS-BDP-FL, Jena Bioscience) to nucleotide-free μOR-G_i_ complexes was measured as previously described (*91*). Briefly, complexes of ligand-bound μOR, unlabeled Gα_i_-Δ6-90/238C and Gβγ were purified by anti-Flag and added at a final concentration of 500nM to a 500 μL quartz cuvette containing 1 μM GTPγS-BDP-FL. Fluorescence was recorded at 512 nm (495 nm excitation) for 500 s at room temperature using a Fluorolog spectrophotometer (HORIBA) with magnetic stirring. All experiments were performed in a buffer containing saturating ligands and 20 mM HEPES pH 7.4, 100 mM NaCl, 0.01% L-MNG, 0.001% CHS, 2mM MgCl_2_ and 100 μM TCEP. Mean association times for GTPγS-BDP-FL binding were calculated from resulting kinetic spectra by fitting a pseudo first-order association function using non-linear regression in Python. We note that the C325A mutation contained in the Δ6 cysteine construct affects the rate of spontaneous GDP release but reportedly not GTP binding^89^.

## Acknowledgements

We thank the Luke Lavis lab for the gift of the JFX673-peg4 conjugate fluorophore; Roger Sunahara and Dirk Siepe for helpful discussions; and Elizabeth White for technical laboratory assistance. Funding support for J.J. was from NIH (K99GM147609) and the Damon Runyon Cancer Research Foundation (DRG-231 8-18). R.V.S. was supported by an Interdisciplinary Fellowship from the Wu-Tsai Institute (Stanford University). S.C. was supported by NIH (R01GM 143554) and the Eleftheria Foundation. B.K.K. was supported by NIH (1R35NS137408).

## Author contributions

JD, DH, JJ, RVS, CS, SC and BKK designed the single molecule experiments. JD, JJ and DH purified proteins, while JD labeled the proteins and prepared complexes for imaging. JD performed nucleotide turnover and binding assays. JD and RVS collected single-molecule data and analyzed it. RVS, JD and SC developed the imaging platform and code. JD, RVS and BKK wrote the manuscript and interpreted the results with input from all authors. RVS and BKK supervised the project.

## Competing interests

BKK is a co-founder and consultant for ConfometRx, Inc. RVS is the director of drug discovery at Greenstone Biosciences.

**Correspondence and requests for materials** should be addressed to Brian Kobilka.

## Data availability statement

The data supporting the findings of this study are available from the corresponding authors after publication. No unique materials or reagents were generated.

## Code availability

Custom MATLAB code used for smFRET data processing is available at: https://github.com/JonathanDeutsch/Freton.

## Extended Data Figures

**Extended Data Fig 1.**
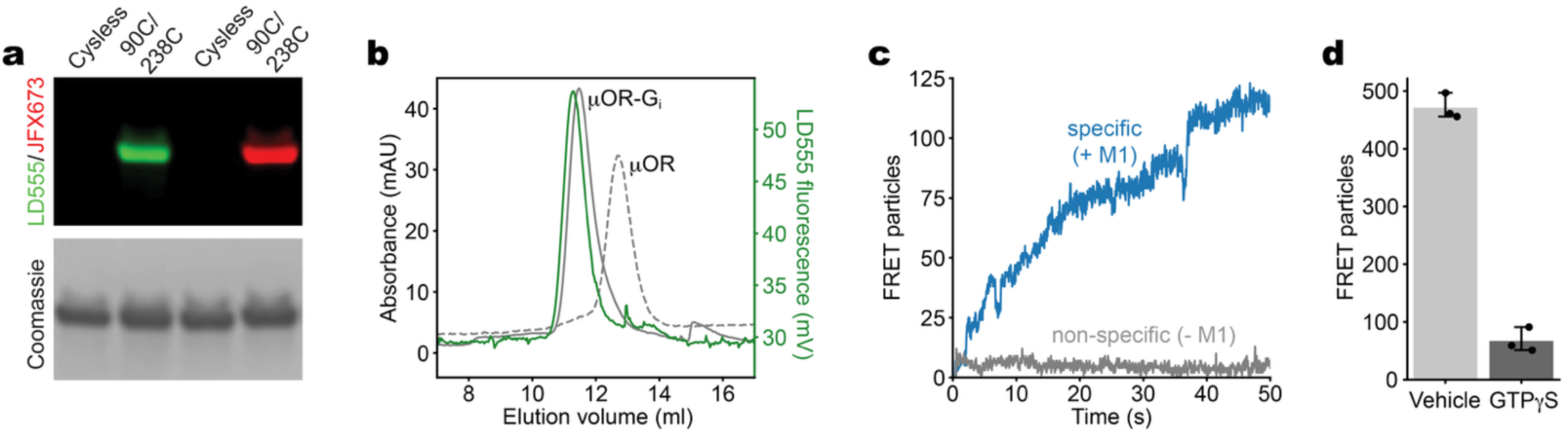
smFRET experimental controls. **a**, Site-specific labeling: SDS-PAGE analysis of purified Gα_i_Δ6 ‘cys-less’ background with and without engineered cysteines (R90C/E238C) labeled with either LD555 (left) or JFX673 (right) visualized using fluorescence imaging (top; 532 nm or 633 nm illumination), with Coomassie staining (bottom) as a gel-loading control. **b**, Representative room-temperature size exclusion chromatography (SEC) profiles of buprenorphine-bound μOR alone (dashed line) and as a ternary complex with dye-labeled G_i_ heterotrimer (absorption in gray; LD555 fluorescence in green). **c**, Specificity of M1 fab-mediated immobilization: Uptake of 200 pM labeled μOR-G_i_ onto passivated quartz slides coated with M1 fab fragment (blue; specific) and without M1 fab (gray; non-specific) recorded using 100 ms TIRF imaging at low laser power (1 mW at the source) to prevent photobleaching. **d**, Histogram of immobilized FRET particles (mean ± 95% c.i., n = 3 movies of the same sample) remaining after injection of buffer (vehicle) or 10 μM GTPγS into the imaging chamber shows GTPγS binding and functional dissociation of μOR-G_i_ on the surface.

**Extended Data Fig 2.**
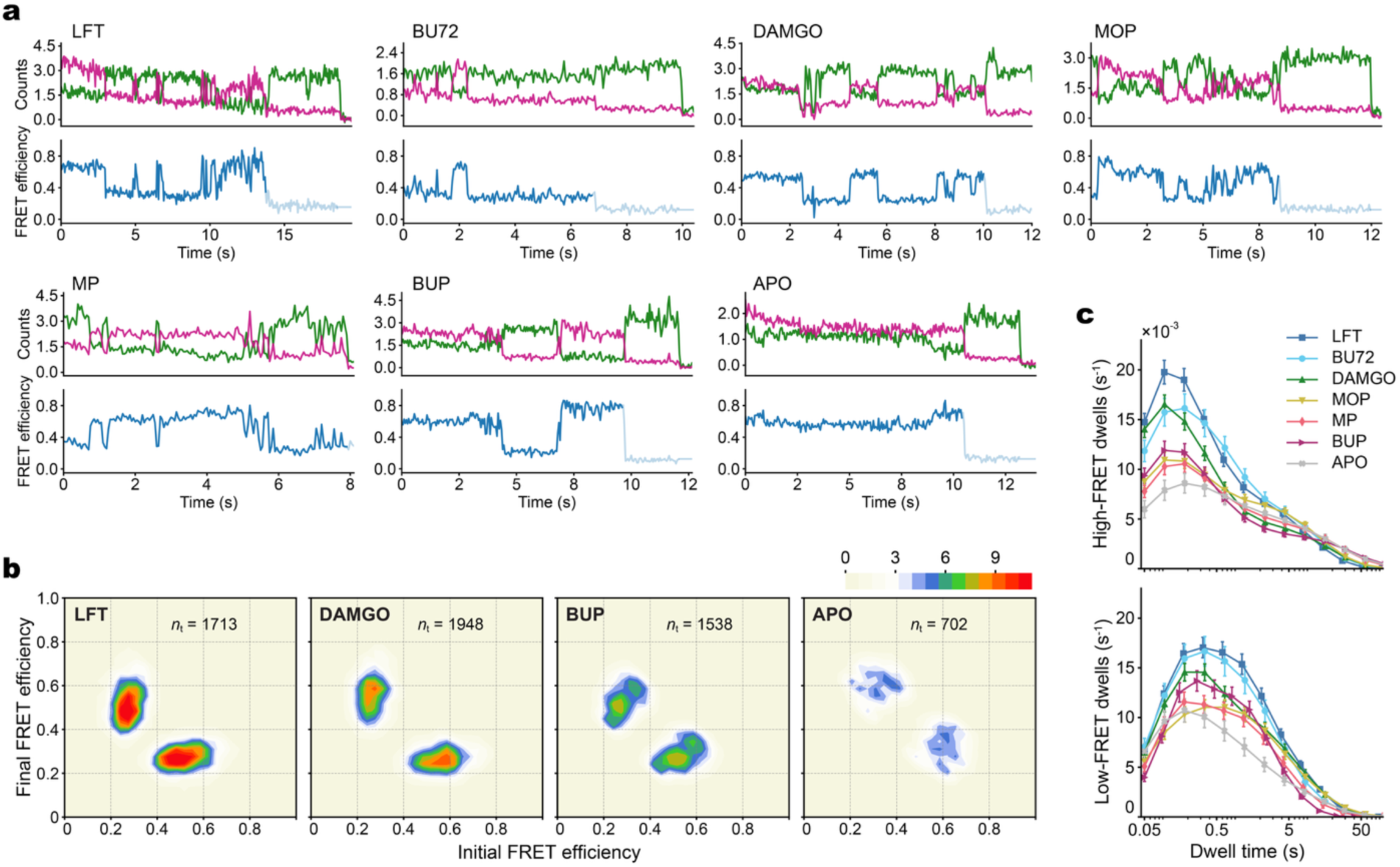
Analysis of AHD opening and closing in μOR-G_i_. **a**, Sample fluorescence (LD555 in green and JFX673 in magenta; units, 10^4^ counts) and FRET efficiency time traces for M1-Fab-immobilized μOR-G_i_ complexes in the indicated ligand and unliganded states. **b**, Transition density plots displaying the mean FRET values before (x-axis) and after (y-axis) each transition (*n*_*t*_ total transitions per condition) for a representative selection of the tested ligands. Scale bar, 10^−3^ transitions per bin per second. **c**, Dwell time distributions of the μOR-G_i_ AHD in the open, low-FRET state (top) and closed, high-FRET state (bottom). Error bars, mean ± s.d. of 100 bootstrap samples of the FRET traces.

**Extended Data Fig 3.**
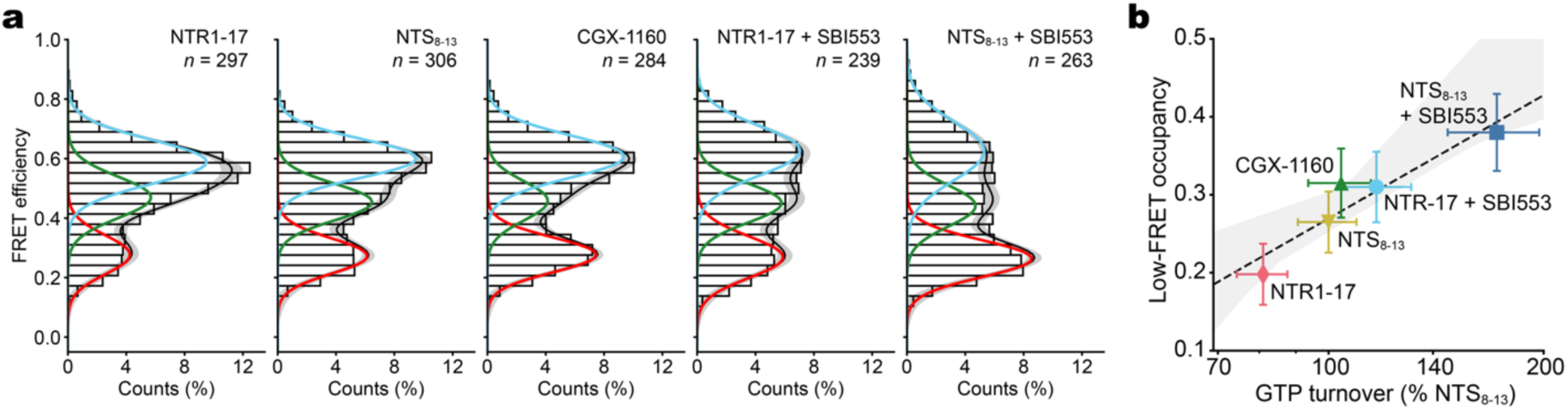
Ligand-induced AHD separation in NTSR1-G_i_. **a**, Population FRET efficiency histograms for surface-tethered NTSR1-G_i_ ternary complexes bound to the indicated ligands, with three-state GMM fits showing the closed (green) and open (cyan and red) states (*n*, number of traces). Error bands, s.d. of the GMM fits from 100 bootstrap samples of the FRET traces. **b**, Correlation scatter plot of ensemble low-FRET, open-state occupancy versus log mean GTP turnover activity (three technical replicates) with R^2^ = 0.89. Error bars, 95% c.i.; error band, bootstrapped 95% c.i. of the regression estimate. Ligands include the full agonist NTS_8-13_, the active fragment of the neurotensin peptide; contulakin-G (CGX-1160), a glycopeptide derivative of NTS found in cone snail toxin; TC-NTR1-17 (NTR1-17), a synthetic partial agonist; SBI-553, an allosteric modulator.

**Extended Data Fig 4.**
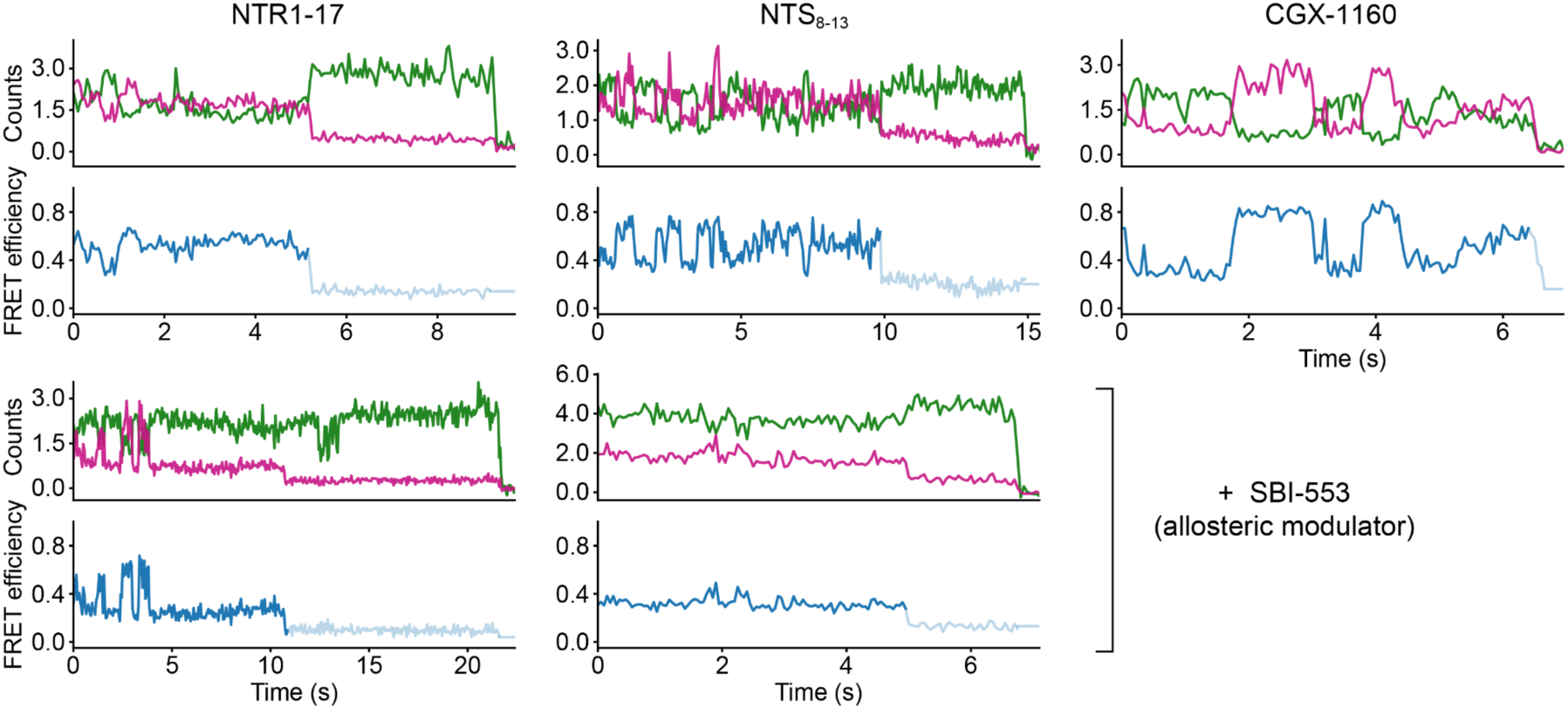
Example smFRET traces of the NTSR1-G_i_ AHD. Example donor and acceptor fluorescence (LD555 in green; JFX673 in magenta) and FRET traces for surface-tethered NTSR1-G_i_ bound to the indicated orthosteric and allosteric ligands.

**Extended Data Fig 5.**
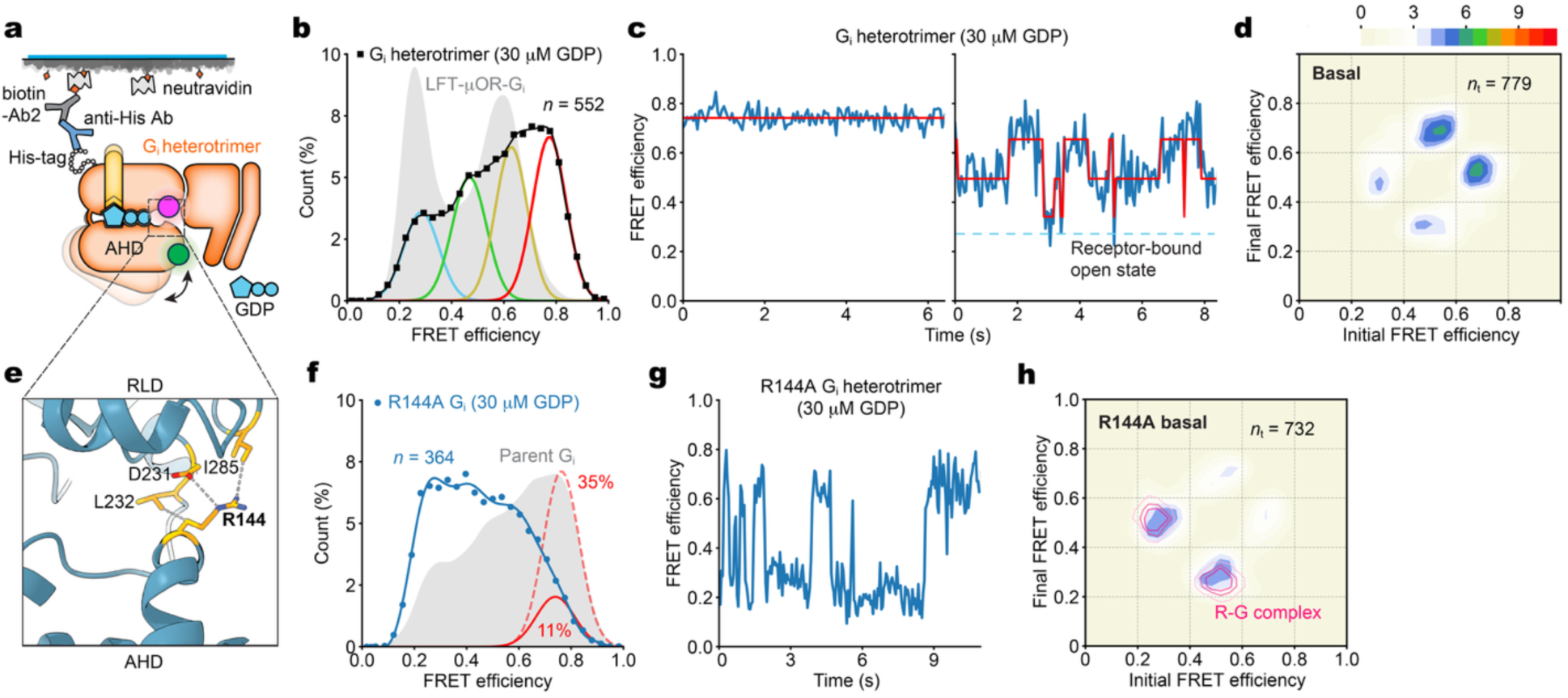
AHD dynamics in the basal state. **a**, Labeled Gα_i_Δ6-90C/238C heterotrimers were tethered to quartz slides via an N-terminal His tag on the Gα subunit and a biotin-mouse antibody bridge. TIRF imaging of the surface-tethered G_i_ AHD, without and with an alanine substitution at R144, was performed in physiologic 30 μM GDP. **b-d**, Population FRET efficiency histogram (squares; *n*, number of traces) with spline and four-state GMM fits (black and colored lines, respectively) (**b**), example FRET and predicted state traces (**c**), and transition density plot for the parent (‘WT’) construct (**d**). **e**, Structure of GDP-bound Gα_i1_ (PDB: 1GP2) showing R144 contacts stabilizing the inactive, closed state at the distal interdomain interface. **f,** Population FRET efficiency histogram (circles) from imaging R144A shows a low-FRET shift relative to WT (red lines are high-FRET gaussians with percent occupancy from four-state fits). **g,h**, Example smFRET trace (**g**) and transition density plot (**h**) showing the effects of R144A on AHD dynamics. Pink lines are overlays of major transition density contours generated by a receptor-G protein (‘R-G’) complex (BU72-μOR-G_i_). Scale bar, 10^−3^ transitions per bin per second (cf. Extended Data Fig. 2b).

**Extended Data Fig 6.**
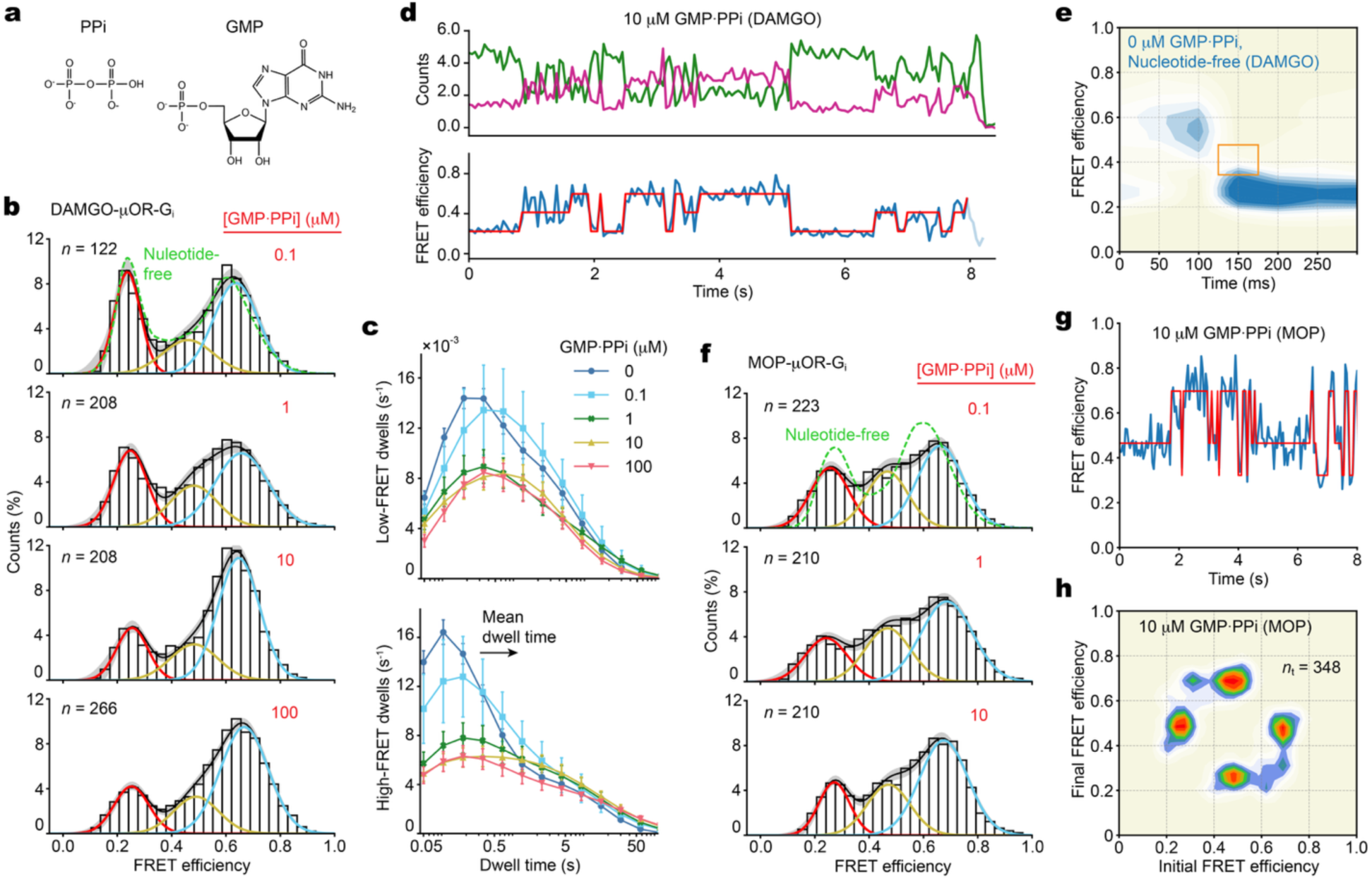
Additional data from imaging μOR-G_i_ with GMP·PPi. Surface-tethered μOR-G_i_ was imaged in a buffer containing various concentrations of equimolar GMP and PP_i_ with either DAMGO- or morphine-occupied receptors. **a**, Structure of PPi and GMP. **b-d**, Population FRET histograms (**b**), dwell time distributions (two state model; see supplementary information) (**c**), and example fluorescence and smFRET traces (**d**; 10 μM GMP·PPi) from experiments imaging DAMGO-μOR-G_i_. **e**, Synchronization plot of high-to-low transitions in the nucleotide-free state for DAMGO-occupied complex (cf. Fig. 3f). **f**, Population FRET histograms from titration experiments with morphine (MOP). **g,h**, smFRET trace (**g**) and transition density plot (**h**) in saturating (10 μM) GMP·PPi from experiments with morphine-bound μOR. Errors are mean ± s.d. of 100 bootstrap samples of the traces.

**Extended Data Fig 7.**
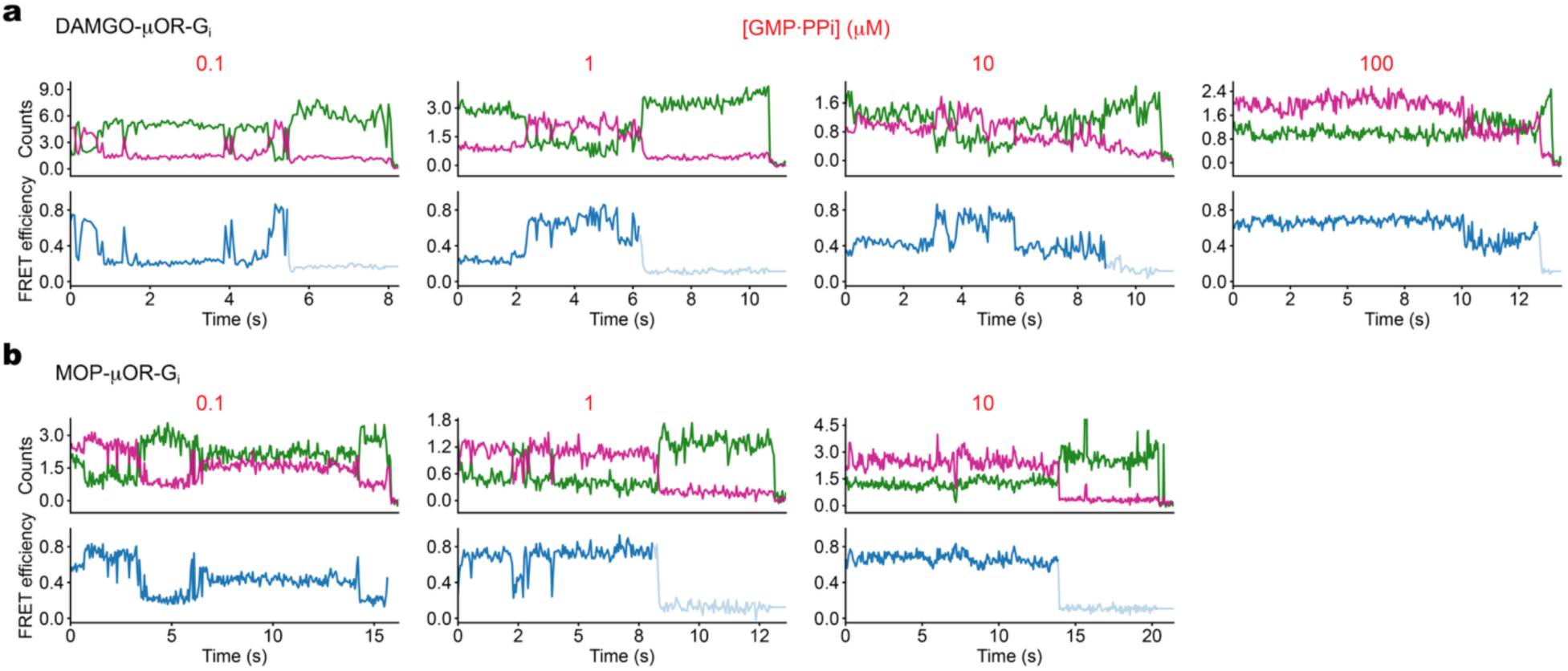
Example smFRET traces with GMP·PPi. **a,b**, Sample trace recordings of the fluorescence (LD555 in green and JFX673 in magenta; units, 10^4^ counts) and FRET efficiency from experiments imaging surface-tethered μOR-G_i_ bound DAMGO (**a**) and MOP (**b**) in buffer containing the indicated concentrations of equimolar GMP and PPi.

